# Lack of Bcl6 promotes anti-tumor immunity by activating glycolysis to rescue CD8 T cell function

**DOI:** 10.1101/2025.04.01.646721

**Authors:** Fangkun Luan, Yunqiao Li, Jia Ning, Jenny Tuyet Tran, Tanya R. Blane, Raag Bhargava, Zhe Huang, Changchun Xiao, David Nemazee

## Abstract

T cells are one of the most powerful weapons to fight cancer; however, T cell exhaustion and dysfunction restrict their long-lasting function in anti-tumor immunity. B-cell lymphoma 6 (BCL6) has many functions in CD8 T cells but it is unclear how it regulates the effector function and exhaustion of CD8 cells. Overall, a low level of BCL6 mRNA in human cancer samples is associated with better outcomes. We found that BCL6 deficiency in activated CD8 T cells enhanced tumor repression in multiple mouse models. More IL-2-expressing CD8 T cells and reduced proportions of exhausted or dysfunctional CD8 T cells were detected within tumors when Bcl6 was knocked out upon T cell activation. Glycolysis was promoted in BCL6-deficient CD8 T cells owing to derepression of glucose transporter GLUT3 (encoded by Slc2a3). The BCL6 inhibitor Fx1 promoted anti-tumor immunity in a T cell-dependent manner. These findings suggest a novel pathway to restore effector function of CD8 T cells by changing their energy utilization pathways to facilitate long-term tumor resistance.

**Summary Blurb:** BCL6 limits CD8 T cell responses to tumors by inhibiting Glut3 expression. Conditional deletion of *Bcl6* in activated CD8 T cells or pharmacological inhibition of BCL6 in mice represses tumor growth.

## Introduction

Upon T cell activation, naïve CD8 T cells differentiate into either cytotoxic effector or memory T cells. Effector T cells rapidly proliferate and secrete killing cytokines to help with pathogen clearance. After the acute response, only a small number of effector T cells survive and develop into memory cells to provide long-term immune protection. When T cells are exposed to persistent antigen stimulation, such as chronic viral infection or tumors, activated T cells become exhausted or dysfunctional, losing their effector functions. Compared to effector or memory T cells, the category of exhausted T cells encompasses a broad definition, characterized by persistently high expression of inhibitory receptors, such as PD1, TIM3, LAG3, CTLA4 and TIGIT, reduced proliferative ability upon stimulation, decreased effector cytokine secretion and a unique epigenetic profile [1, 2]. Exhausted T cells appear to lose their ability to kill target cells and occupy a stable T differentiation state [3] distinct from memory T cells. Moreover, the exhausted T cell population is heterogeneous, containing at least progenitor and terminal subsets [4]. Progenitor-exhausted T cells have high levels of expression of TCF1, SLAMF6 and CXCR5 [5–8], whereas cells expressing TIM-3 and CD39 identify terminally exhausted T cells [9, 10]. Markers Ly108 (encoded by *Slamf6*) and CD69 can further divide four subsets among those exhausted CD8 T cells [11]. In addition to these characteristics of exhausted T cells, their energy metabolism is different from effector or memory T cells. Effector T cells rely on aerobic glycolysis for rapid proliferation and effector activities, while memory T cells perform elevated OXPHOS and autophagy to maintain their long-term survival and function [10, 12–15]. In contrast to the energy requirements of effector and memory T cells, exhausted T cells demonstrate bioenergetic insufficiencies, such as suppressed mitochondrial respiration and poor glycolysis [16–18].

BCL6 is a transcriptional repressor reported to participate in many types of regulation of T and B cells [19]. BCL6 binds to the promoter region of *Gzmb* (encoding Granzyme B) to repress its expression in naïve CD8 T cells, rather than in activated CD8 T cells [20]. BCL6 is also critical for maintaining and generating memory CD8 T cells [21]. The expression level of BCL6 exhibits a strong positive correlation with the number of central memory CD8 T cells (TCM) upon antigen stimulation and contributes to antigen-specific TCM secondary expansion [22]. As a well-established antagonist of BLIMP1 activity, BCL6 promotes a stem-like CD8+ T cell formation with memory potential and high proliferation capacity [23]. Since memory T cells or exhausted T cells are generated from effectors, it is unclear if BCL6 has any role in the differentiation of T cell exhaustion, its functional window period or whether it can be used as a target of cancer treatment.

## Methods

### Mice and Tumor Models

*Bcl6*^fl/fl^ conditional knockout mice (*Bcl6*^fl/fl^ CD4-cre, *Bcl6*^fl/fl^ Mb1-cre, *Bcl6*^fl/fl^ Ox40-cre and *Bcl6*^fl/fl^ Gzmb-cre) were generated and maintained in-house. Wild-type C57BL/6 and *Rag1*^−/–^ (*Rag1*^tm1Mom^) mice were obtained from Jackson Labs. Mice were used at 6–8 weeks of age, and all procedures adhered to institutional animal care guidelines. Subcutaneous tumor models were established by inoculating 1 × 106 MC38, LLC1, or B16F10 cells per mouse. Tumor growth was monitored every three days using digital calipers.

### Reagents and Antibodies

All cell culture experiments utilized DMEM (Fisher Scientific) supplemented with 10% fetal bovine serum (Gibco). Fluorescently conjugated antibodies for flow cytometry were purchased from BD Biosciences. The Cut&Run Assay Kit was obtained from Vazyme, and RNA sequencing libraries were prepared using the SMART-Seq v4 Ultra Low Input RNA Kit (Takara Bio). For qPCR, reverse transcription was performed using HiScript III RT SuperMix (Vazyme), and real-time PCR utilized qPCR SYBR Green Master Mix (Vazyme). Fx1 (Selleckchem) was dissolved in a vehicle containing 40% DMSO, 40% PEG3000, and 20% PBS, and administered intraperitoneally at 20 mg/kg.

### Flow Cytometric Analysis

Tumor-infiltrating lymphocytes (TILs) were isolated by enzymatic digestion of tumor tissue with collagenase I (1 mg/mL) and DNase I (0.1 mg/mL). Cells were stained in FACS buffer (PBS supplemented with 2% FBS) and analyzed on a BD LSRFortessa flow cytometer. Data were processed using FlowJo software, and gates were set based on fluorescence-minus-one (FMO) controls.

### RNA Sequencing

CD8+PD1+ TILs were sorted to >95% purity. RNA was extracted and libraries were generated using the SMART-Seq v4 Kit. Sequencing was conducted on an Illumina NovaSeq platform, with differential gene expression analyzed using DESeq2. Enrichment analyses were performed with the KEGG pathway database.

### Metabolic Assays

Extracellular acidification rates (ECARs) were measured using the Seahorse XF Glycolysis Stress Test Kit (Agilent) on an XF96 Analyzer. For glucose uptake assays, cells were cultured in glucose-free medium and incubated following CD3/CD28 stimulation.

### Chromatin Immunoprecipitation (Cut&Run)

CD8 T cells were isolated from the spleen of WT mice using EasySep™ Mouse CD8+ T Cell Isolation Kit (StemCell Technologies), and their lysates for ChIP were assayed directly. Cut&Run assays were performed as described by the manufacturer of Hyperactive pG-MNase CUT&RUN Assay Kit (Vazyme). Anti-BCL6 (Cell Signaling Technology) and isotype control antibodies (Cell Signaling Technology) enriched DNA-protein complexes. qPCR was conducted to assess binding at the *Slc2a3* promoter region, with enrichment quantified relative to IgG controls. Primers for qPCR: Forward GGAGCGGTGAAGATCAGATAAG; Reverse GAAGCCCAGCCTACCTATTT.

### Luciferase

On the day of transfection, EL4 suspension cells were seeded 5×10^4^/well in a 96-well plate (clear bottom) first. To a sterile tube, Opti-MEM medium was added to make the final volume 50 μl after adding the transfection-grade DNA (10 μl/well). 200 ng, 100 ng, 50 ng, or 25 ng of pCMV-2b-mBCL6-FL plasmid or an equal amount of the empty vector was added into the medium followed by 250 ng reporter plasmid pGL4.10[Luc2]-Slc2a3(−127,+129) and 50 ng internal control vector pGL4.74[hRluc/TK]. For a 3:1 ViaFectTM transfection reagent (Promega): DNA ratio, we added 1.5 μl of the reagent to the media, mixed immediately then waited for 10 min at room temperature. To each well 10 μl of the mixture was added to the plate, with gentle pipetting, then the plate was returned to the incubator. After 48 hours, we followed the instructions of the Dual-Glo Luciferase assay system (Promega) to capture the firefly and Renilla Luminescence in a luminometer, and calculate the fold changes.

## Statistical Analysis

Data are presented as mean ± SEM. Statistical significance was determined using GraphPad Prism 9.0, employing unpaired two-tailed t-tests or two-way ANOVA with multiple comparison corrections where applicable. Statistical thresholds for significance were set at p < 0.05.

## Results

### Conditional knockout of *Bcl6* in activated CD8 T cells represses tumor growth

To investigate the association between *Bcl6* expression and the outcomes of tumor samples, we took advantage of the GEPIA database [24], which can be used to analyze the RNA sequence data of tumor samples under the TCGA and GTEx projects. We found that tumor cases with lower levels of *Bcl6* mRNA had favorable overall survival rates in all types of cancer (Fig. 1A) and this phenotype was confirmed in colon adenocarcinoma (COAD, with p=0.049), pancreatic adenocarcinoma (PAAD, with p=0.021) and, possibly, rectum adenocarcinoma (READ, no significant difference but trend was shown) (Fig. S1A-C). Moreover, tumor-infiltrating CD8 T cell levels and *Bcl6* expression showed a positive correlation in COAD, PAAD and READ samples (Fig. S1D-F), suggesting that BCL6 promotes CD8 T cell proliferation within tumors while suppressing their killing function.

**Figure 1.**
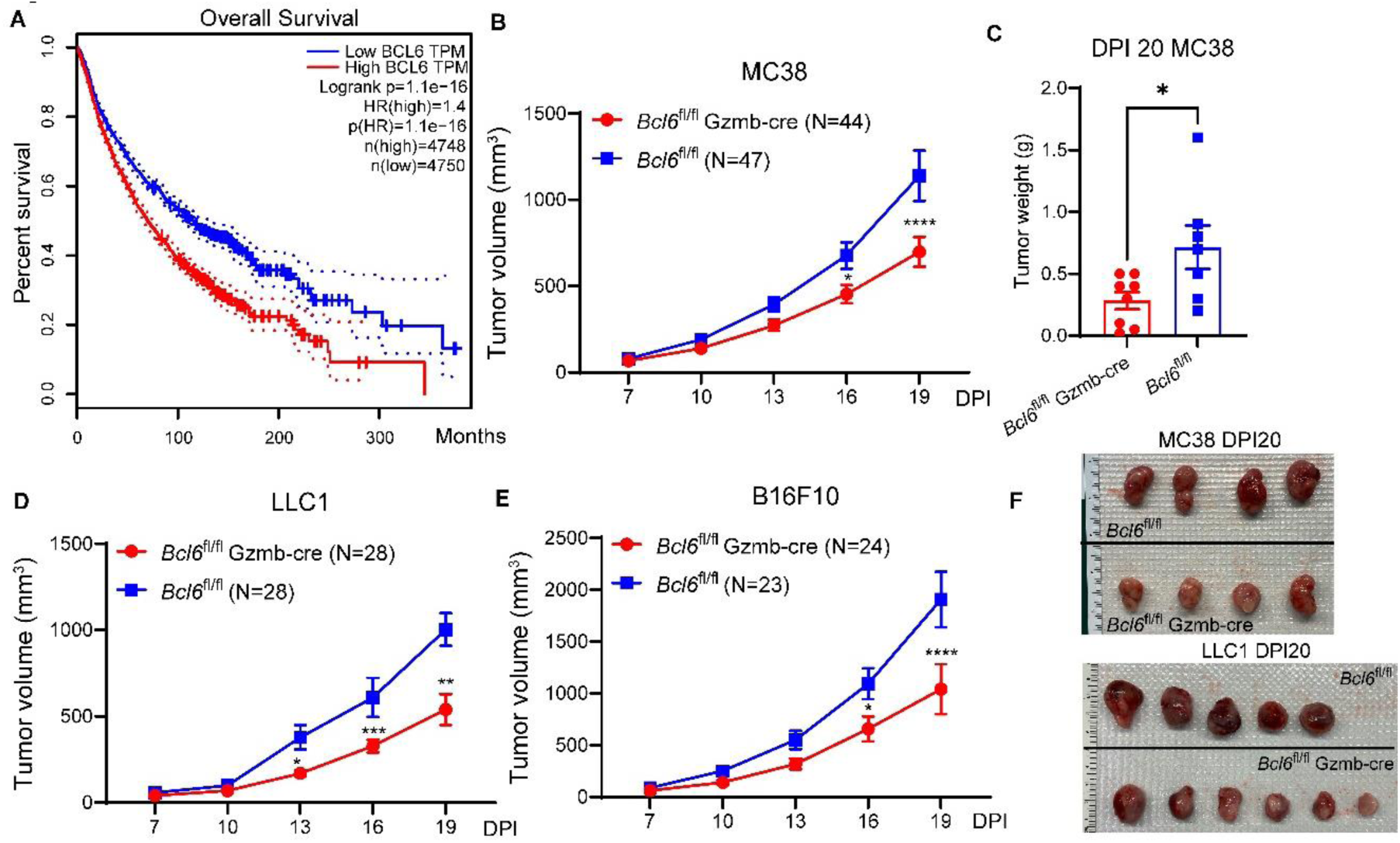
Low level of BCL6 is correlated with improved survival and reduced tumor growth. **(A)** Relationship between BCL6 expression and survival of patients with all types of cancer (http://gepia.cancer-pku.cn/index.html). **(B-C)** *Bcl6*^fl/fl^ and *Bcl6*^fl/fl^ GzmB-cre mice were inoculated with 1×10^6^ MC38 tumor cells on day 0. Tumors were measured every 3 days from day 7 post-inoculation (DPI) revealing slower tumor growth in *Bcl6*^fl/fl^ GzmB-cre mice (B). Comparison of tumor weights on DPI 20 (C). **(D-E)** Similar tumor repression phenotypes were observed in *Bcl6*^fl/fl^ GzmB-cre mice against LLC1 (D) and B16F10 (E) tumor cell inoculations. (F) Visual analysis of tumors of MC38 (upper) and LLC1 (lower) on DPI20 comparing *Bcl6*^fl/fl^ GzmB-cre to *Bcl6*^fl/fl^ tumor recipients. Tumor growth data were pooled from at least 4 independent experiments. Each point in graph (C) represents the value obtained in an independent mouse. Bars are presented as mean ± SEM. P values were calculated with the two-tailed unpaired t-test (C) or two-way ANOVA (B, D, E). * denotes P < 0.05.

To study the function of BCL6 in the tumor immune response, we analyzed tumor growth in various conditionally deficient BCL6 models. *Bcl6*^fl/fl^ CD4-cre mice, which lack BCL6 in both CD4 and CD8 T cells, starting at the double positive stage of T cell development, showed inhibited growth of MC38, LLC1 (Lewis lung carcinoma) and B16F10 (melanoma) tumors compared to *Bcl6*^fl/fl^ control mice (Fig. S2A-G). This contribution to tumor resistance likely did not come from CD4 T cells as *Bcl6*^fl/fl^ OX40-cre mice showed tumor growth curves comparable to *Bcl6*^fl/fl^ controls (Fig. S2H). In contrast, *Bcl6*^fl/fl^ Gzmb-cre mice showed repressed growth of MC38, LLC1 and B16F10 tumors, suggesting that the lack of BCL6 in activated CD8 T cells was crucial for enhanced anti-tumor immunity (Fig. 1B-F).

### Lack of BCL6 retards exhaustion of CD8+ T cells and restores their effector functions

The data described above, indicating that repressing BCL6 expression in activated CD8 T cells can benefit tumor control and may result in changes in CD8 T cell effector function, encouraged us to compare MC38 tumor-infiltrating CD8 cells in *Bcl6*^fl/fl^ and *Bcl6*^fl/fl^ Gzmb-cre mice. On day 8 post-inoculation (DPI 8) of MC38, *Bcl6*^fl/fl^ Gzmb-cre mice had more tumor-infiltrating, effector IFNγ+ CD8 T cells compared to *Bcl6*^fl/fl^ mice, though significantly reduced proportions of IFNγ+ CD8 T cells were observed in both groups by DPI 20 (Fig. 2A). However, the proportions of tumor-infiltrating cells producing IL-2 increased in BCL6-deficient CD8 T cells on both DPI 8 and DPI 20 (Fig. 2B). There was no difference found in TCF1 expression when comparing activated CD44+ CD8 T cells of *Bcl6*^fl/fl^ Gzmb-cre and *Bcl6*^fl/fl^ mice (Fig. 2C), suggesting that knocking out BCL6 during CD8 T cell activation may not have influenced their stemness and memory differentiation. On the other hand, the numbers of tumor-infiltrating “exhausted” TIM3+PD1+ CD8 T cells were significantly reduced in *Bcl6*^fl/fl^ Gzmb-cre mice on DPI 20 (Fig. 2D). Moreover, the CD39+PD1+ and CD69+PD1+ dysfunctional CD8 T cells, which were obviously increased in proportion within tumors from DPI 8 to DPI 20 in control mice, were significantly reduced on DPI 20 in *Bcl6*^fl/fl^ Gzmb-cre mice (Fig. 2E, F). These data suggest that as the size of inoculated tumors increase the function of CD8 T cells transforms from effector to exhaustion or dysfunction, and that the elimination of BCL6 in activated CD8 T cells can reverse this process to some extent, allowing them to exert their killing effector for a longer time.

**Figure 2.**
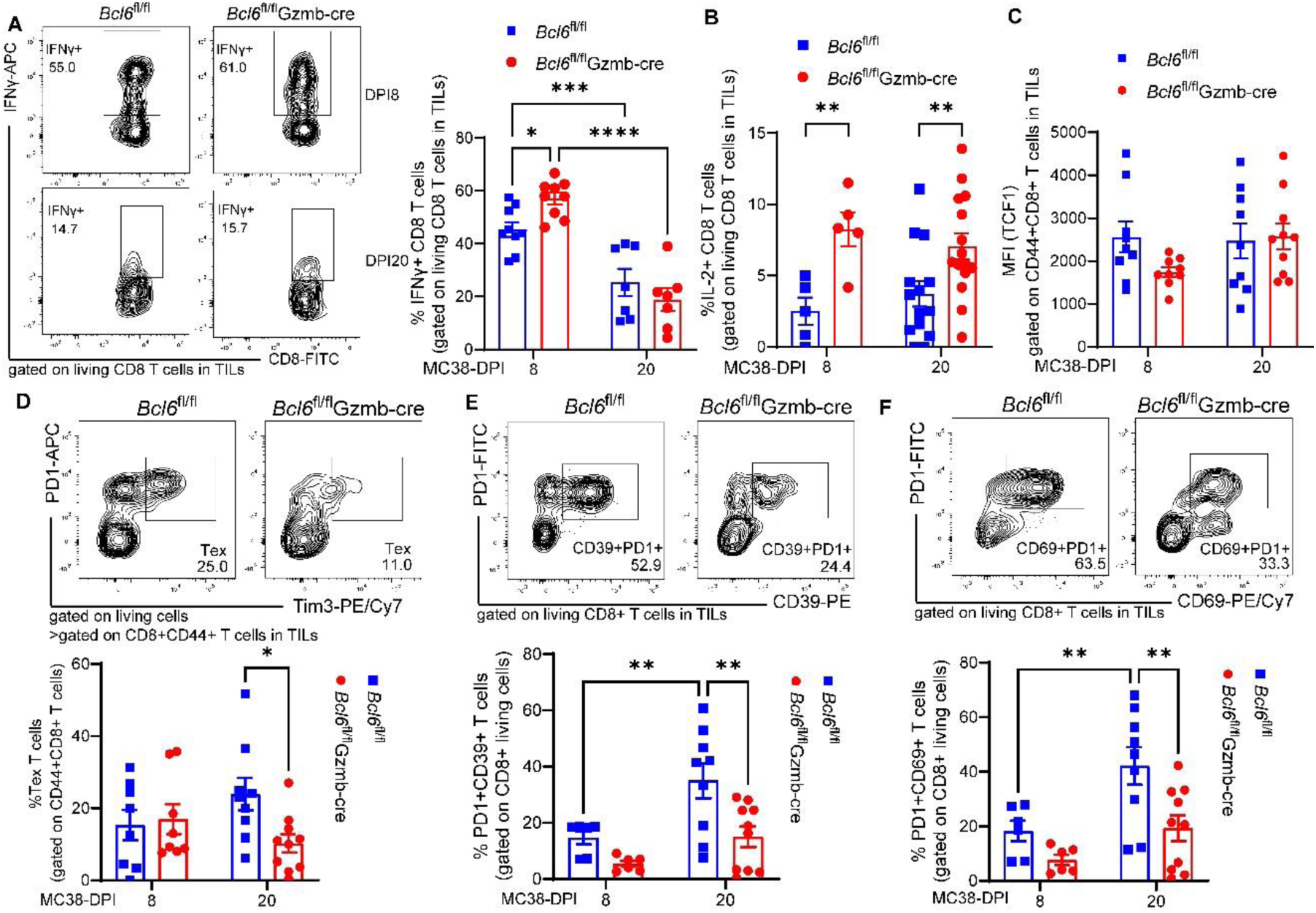
T cell dysfunction is lessened in BCL6-deficient CD8 T cells. **(A)** Representative flow cytometry plots and frequencies of MC38 tumor-infiltrating IFNγ+ CD8 T cells on DPI8 (upper and left panel) and DPI20 (lower and right panel). **(B)** Frequency of IL-2+ CD8 T cells within tumors on DPI8 (left) and DPI20 (right). **(C)** Levels of TCF1 expression on tumor-infiltrating CD44+CD8+ T cells on DPI8 (left) and DPI20 (right). **(D-F)** Representative flow cytometry plots in the upper panel exhibited the following categories of CD8 T cells on DPI20: (D) tumor-infiltrating exhausted, PD1+Tim-3+; (E) dysfunctional, PD1+CD39+; and (F) PD1+CD69+ T cells. Lower panels show frequencies of these subpopulations on DPI8 (left) and DPI20 (right). Flow cytometry data were pooled from at least 3 independent experiments. Each point in graphs (A-F) represents the value obtained in an independent mouse. Bars are presented as mean ± SEM. P values were calculated with the two-tailed unpaired t-test (B), Two-way ANOVA (A, D-F). *, **, ***, **** denote P < 0.05, 0.01, 0.001, 0.0001, respectively.

### Glycolysis is improved in BCL6-deficient CD8 T cells

To investigate how BCL6 regulates activated CD8 T cell function, we performed bulk RNA-seq, comparing sorted living PD1+CD8 T cells isolated from DPI 10 tumors of MC38-inoculated *Bcl6*^fl/fl^ Gzmb-cre (BCL6-KO) to those of *Bcl6*^fl/fl^ (BCL6-WT) mice. EnrichKEGG analysis of the upregulated genes indicated that cell cycle, biosynthesis and energy metabolism regulation were stimulated in BCL6-KO CD8 T cells (Fig. 3A, Supplementary Table 1). Consistently, the KEGG enrichment network showed improvement of glycolysis when BCL6 expression was restricted (Fig. 3B, Supplementary Table 1). Genes associated with glycolysis were increased in activated BCL6-KO CD8 T cells (Fig. 3C). To test glycolysis function, we performed metabolic stress tests using naive or TCR-stimulated CD8 T cells from *Bcl6*^fl/fl^ CD4-cre mice. Basal glycolysis in BCL6-KO CD8 T cells was significantly higher than in BCL6-WT CD8 T cells, and the activated BCL6-KO CD8 T cells had better glycolytic compensation when mitochondrial respiration was inhibited (Fig. 3D). Glucose uptake of CD8 T cells was detected with or without CD3/CD28 stimulation conditions; notably, BCL6-KO CD8 T cells had significantly enhanced glucose uptake upon TCR stimulation compared to BCL6-WT CD8 T cells (Fig. 3E). Higher levels of glucose transporter GLUT3 were observed in both resting and activated BCL6-KO CD8 T cells (Fig. 3F), consistent with their stronger glucose uptake. Increased expression of GLUT3 was also detected on tumor-infiltrating CD8+PD1+ T cells of *Bcl6*^fl/fl^ Gzmb-cre mice on DPI 20 (Fig. 3G). These data demonstrate enhanced glycolysis in BCL6-deficient CD8 T cells through up-regulation of glucose transporter GLUT3 expression and GLUT3-mediated glucose uptake.

**Figure 3.**
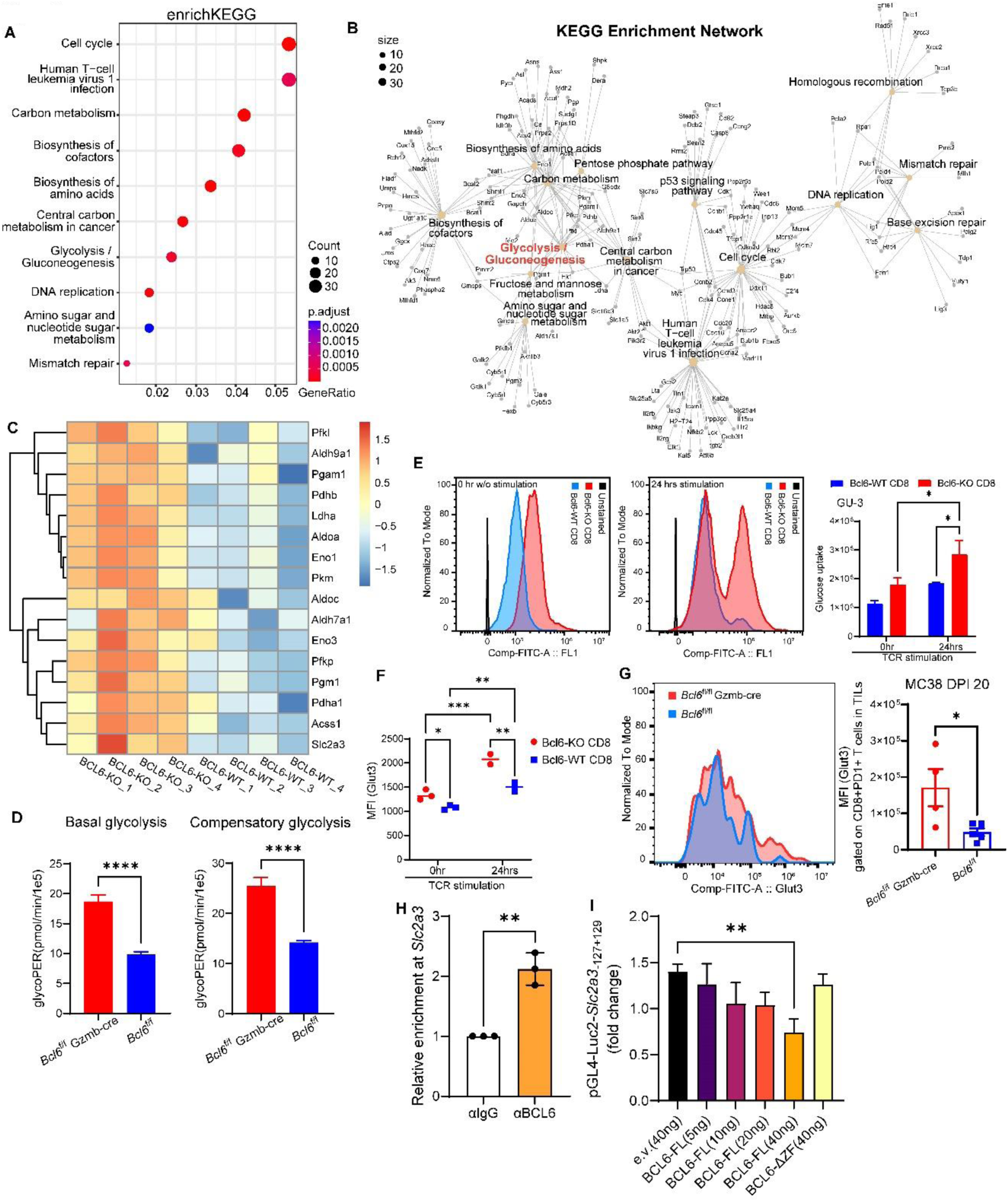
GLUT3 is repressed by BCL6, in turn regulating CD8 T cell glycolysis. **(A-C)** RNA analysis of CD8 T cells isolated from tumors of MC38-inoculated *Bcl6*^fl/fl^ Gzmb-cre or *Bcl6*^fl/fl^ mice. (A) Dot plots and (B) enrichment network of the enrichKEGG analysis for upregulated genes in activated BCL6-knockout (BCL6-KO) CD8 T cells. (C) Heatmap shows the frequencies of remarkable genes in the glycolytic pathway of individual mice from the BCL6-KO and BCL6-WT activated CD8 T cell groups. **(D-F)** Metabolic analysis of CD8 T cells. (D) Analysis of basal (left) and compensatory (right) glycolysis levels of BCL6-KO vs BCL6-WT CD8+ T cells isolated from *Bcl6*^fl/fl^ Gzmb-cre or *Bcl6*^fl/fl^ mice that had been previously activated on anti-CD3/anti-CD28 plates overnight then stimulated for 48 h with IL-2 and anti-CD28. (E-F) Comparison of glucose uptake and GLUT3 expression in *Bcl6*^fl/fl^ CD4-cre CD8+T cells (BCL6-KO) vs *Bcl6*^fl/fl^ CD8+T cells (BCL6-WT). (E) Elevated glucose uptake in BCL6-KO compared to BCL6-WT CD8 T cells after 24 hrs TCR stimulation. Similar differences were reproduced in a second experiment. (F) Levels of GLUT3 expression on BCL6-KO and BCL6-WT CD8 T cells before and after TCR stimulation. Data were pooled from 2-3 independent experiments. **(G)** Increased GLUT3 detected on tumor-infiltrating PD1+ CD8 T cells in *Bcl6*^fl/fl^ Gzmb-cre mice on DPI 20. Each point represents the value obtained in an independent mouse. **(H)** Analysis of BCL6 binding to the promoter regions of *Slc2a3*, which encodes GLUT3 in mice. Binding activity was confirmed three times independently using CD8 T cells isolated from spleen of WT mice. **(I)** Luciferase assay showing that full-length (FL) BCL6 repressed the expression of the reporter gene (luc2) driven by the *Slc2a3* promoter, and that the inhibition of luc2 signal was lost when BCL6 lacked its DNA binding activity (BCL6-ΔZF). Data were pooled from 4 independent experiments. Bars show mean ± SEM. P values were calculated with the two-tailed unpaired t-test (D, G, H), Two-way ANOVA (E, F, I). *, **, ***, **** denote P < 0.05, 0.01, 0.001, 0.0001, respectively.

### BCL6 binds the promoter region of *Slc2a3* and represses its expression

GLUT3 is encoded by *Slc2a3*. We performed a Cut&Run experiment to quantitate immunoprecipitation of DNA segments bound by transcription factors. Compared to the IgG control group, BCL6 was significantly enriched in the promoter regions of *Slc2a3* (Fig. 3H). Luciferase assay showed *Slc2a3* promoter-driven reporter (Luc2) gene expression was repressed by full-length BCL6, while the suppression of Luc2 expression was recovered when BCL6 lost its DNA binding ability (deletion of ZF fragment, ΔZF [25]) (Fig. 3I). These data indicate that GLUT3 expression is repressed by BCL6 binding to its promoter regions.

### Fx1, a BCL6 inhibitor, suppresses tumors in a T cell-dependent manner

To determine if CD8 T cell exhaustion could be treated pharmacologically, we decided to test the anti-tumor efficacy and mechanism of action of the BCL6 inhibitor Fx1. We first confirmed that BCL6 expression was strongly induced upon T cell activation (Fig. 4A). Fx1 treatment, given by daily intraperitoneal (i.p.) injection starting on day 10 after MC38 or B16F10 tumor inoculation, repressed tumor growth in C57BL/6 mice (Fig. 4B, C). As BCL6 is a master regulator of the cell cycle, we considered the possibility that Fx1 may work to inhibit tumor expansion directly. This seemed unlikely, however, as the anti-tumor efficacy was lost when the therapy was introduced in tumor-bearing *Rag1*^−/–^ mice, which lack T and B cells (Fig. 4B, C). As *Rag1*^−/–^ mice are sufficient in NK cells, the result also indicated that inhibiting BCL6 in NK cells was not as important as targeting BCL6 in T cells or B cells. Moreover, as MC38 and B16F10 growth curves were comparable in *Bcl6*^fl/fl^ Mb1-cre and *Bcl6*^fl/fl^ mice, BCL6 insufficiency or derepression of its antagonized targets, such as BLIMP in B cells, likely does not affect tumor immunity (Fig. 4D, E). Collectively, these data demonstrate that the BCL6 inhibitor Fx1 controlled tumor progression in a T cell-dependent manner. Lack of BCL6 in activated CD8 T cells stimulated the GLUT3-mediated glycolysis pathway, ultimately retarding CD8+ T cell exhaustion and restoring their effector functions. This mechanism explains the T cell dependence of Fx1 drug-induced tumor resistance.

**Figure 4.**
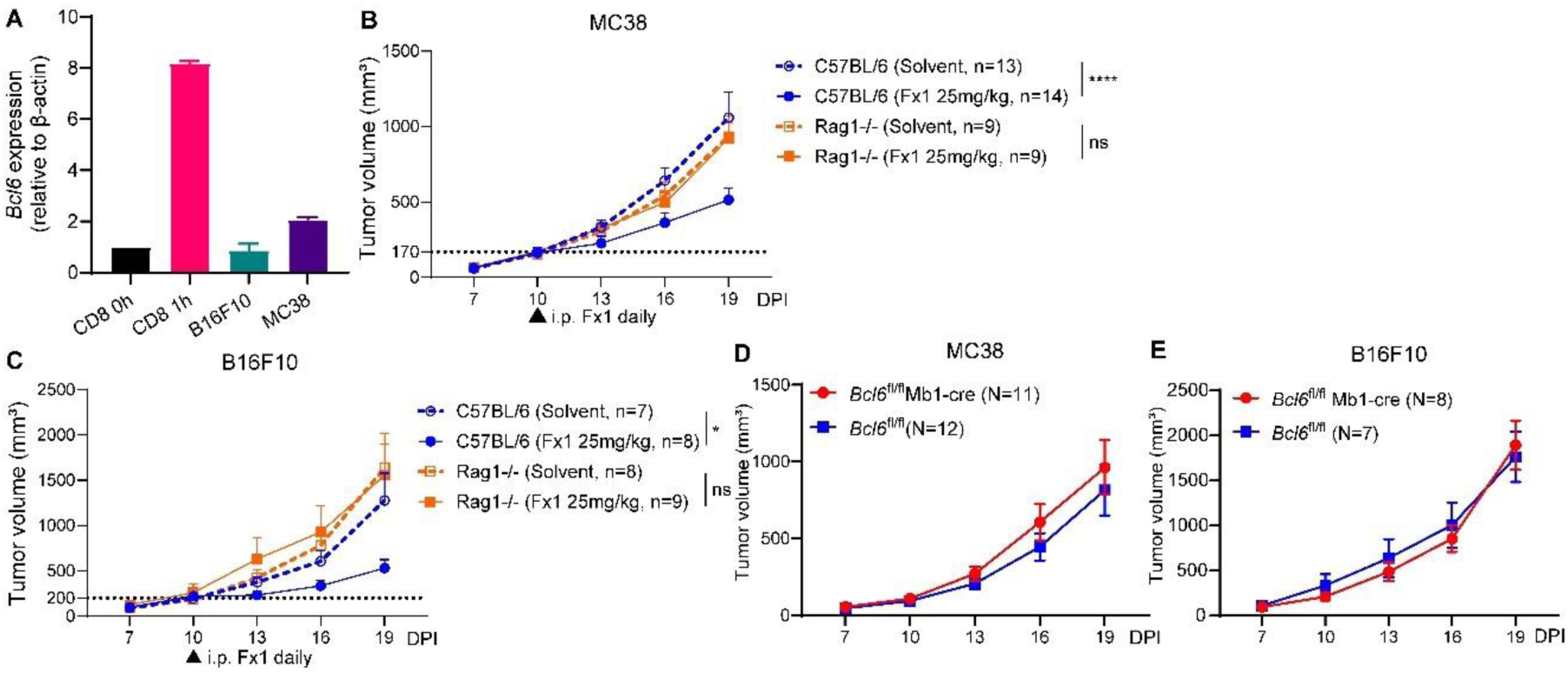
The anti-tumor efficacy of the BCL6 inhibitor Fx1 depends on T cells. **(A)** The relative expression of BCL6 in B16F10 and MC38 tumor cells is comparable to the level in CD8 T cells without TCR stimulation, and BCL6 is quickly increased in CD8 T cells after activation. **(B-C)** Analysis of tumor growth in C57BL/6 and *Rag1*^−/–^ mice implanted with MC38 (B) or B16F10 (C) tumor cells and receiving Fx1 treatment i.p. daily starting from DPI 10. **(D-E)** Analysis of tumor growth of MC38 (D) and B16F10 (E) in *Bcl6*^fl/fl^ Mb1-cre and control *Bcl6*^fl/fl^ mice. Bars show mean ± SEM. P values were calculated with the Two-way ANOVA (B-C). *, **** denote P < 0.05, 0.0001, respectively, and ns denotes P > 0.05.

## Discussion

T cell responses are self-limiting, which may be adaptive late in infection when microbial clearance allows cells to recover from exhaustion. However, in the context of antigen persistence, such as in cancer, prevention or reversal of exhaustion is often desirable. Here we show that genetic or pharmacological suppression of BCL6 in activated CD8 cells promotes tumor control. This biological effect is accompanied by enhanced T cell secretion of functionally important cytokines, such as IFNγ, IL-2, and a reduction in frequency of cells with markers of T cell exhaustion. The mechanism by which BCL6 normally limits the response appears to be in part through restricting glucose transport and glycolysis in CD8 T cells. Our study supports the observation from the study of Chen Dong and colleagues that BCL6 limits T cell function against cancer [23]. However, our mechanistic data provide a distinct interpretation. We show that BCL6 works by directly suppressing a glycolysis stimulation pathway to remodel the effector T cell function late after the activation of CD8 T cells and that introducing a powerful small molecular drug treatment to suppress BCL6 works even after the tumor grows to nearly 200 mm^3^ volume. Furthermore, depletion of BCL6 during activation of CD8 T cells did not result in any difference in TCF1 expression in CD8 T cells, suggesting that disruption of BCL6 during this time window may not induce stem-like/progenitor exhausted T cells at all, thereby inhibiting the generation of terminally exhausted T cells by bypassing the precursor formation pathway. On the other hand, the model that we used, *Bcl6*^fl/fl^ Gzmb-cre, interrupted the powerful function of BCL6 in a narrow cohort of cells in a limited developmental window, leaving other CD8 T cells untouched, and thereby limiting the possibility of affecting cytokine production.

Understanding the mechanisms of T cell exhaustion should facilitate the identification of pharmacological remedies able to restore the cytotoxicity of dysfunctional T cells to control tumor progression. T-cell exhaustion is complex, difficult-to-manipulate, and still not fully understood. Exhausted T cells cannot be simply identified by their expression of inhibitory receptors, however. Many T cell effectors secrete cytolytic cytokines while maintaining high levels of inhibitory molecules [26]. For example, Tc17 cells express transcription factors STAT3 and RORγt, enabling their effector functions by producing IL-17, IL-22, TNF, etc. [26–28]. At the same time, Tc17 cells are increased in Crohn’s disease and highlighted by expression of CD6^high^, CD39, CD69, PD-1, and CD27^low^, which can be targeted to inhibit disease [29]. Metabolomics and mapping of cytokine secretions of different CD8 T cell subtypes may improve our understanding of the development and mechanisms of T cell exhaustion.

Metabolic reprogramming is essential for the activation and function of effector T cells [14, 30, 31]. Upon TCR-mediated activation, T cells increase the expression of glucose transporters and glycolysis enzymes to promote aerobic glycolysis by rapidly stimulating the PI3K/AKT/mTOR1 pathway [32]. However, during continuous antigen stimulation exposure to activated T cells, metabolic alterations of glucose, amino acids and fatty acids are significant features as cells progress from effector T cells to exhausted T cells [30, 32–35]. Shifting from glycolysis to fatty acid oxidation (FAO) in T cells occurs even before T cell dysfunction occurs, suggesting that the metabolic defect is a trigger rather than a consequence of T cell dysfunction or exhaustion [17, 36, 37]. In this study, we found that the inhibition of BCL6 in activated CD8 T cells increased the expression of glucose transporter GLUT3, thereby maintaining glycolysis, and indicating that reversing T cell exhaustion in tumor conditions is a plausible therapeutic strategy. Currently, the most effective immunotherapy against tumors is immune checkpoint blockade (ICB). Targeting exhausted T cells to restore their effector function is an independent, non-mutually exclusive approach clearly worth further investigations.

Glucose transporter family members (GLUT), are known to have distinct sequences, substrate affinity, and kinetic properties that reflect specific roles of glucose homeostasis in different cell types [38]. In CD4 T cells, GLUT1 is induced upon stimulation and is maintained most predominantly, whereas GLUT3 levels become less prominent, and although GLUT6 also increases, its levels are lower than GLUT1[39]. GLUT1 was reported to be the primary high affinity glucose transporter of T cells [40]. However, compared to activated CD4 effectors, CD8+ effector T cells are less dependent on GLUT1 and oxygen levels [41]. It is reported that the normal function of CD8 effector T cells may utilize GLUT3 or GLUT6 expression, since the deletion of GLUT1 on CD8 T cells does not result in their inability to differentiate into effector functions and release cytokines such as GzmB, TNFα, IL-2, and IFNγ [39]. GLUT3 has a higher affinity and increased transport ability for glucose compared to GLUT1 [42]. Overexpression of GLUT3 in murine CD8 T cells promotes glucose uptake and enhances glycogen and fatty acid storage, sustaining effector function [43]. Forced expression of GLUT3 along with tumor-specific chimeric antigen or T-cell receptors in engineered human T cells increases their antitumor ability in the context of adoptive T-cell therapies [44]. Compared to GLUT1, GLUT3 is induced primarily in the IL-2 signaling branch [45], which also supports our observations of higher levels of IL-2 on BCL6-deleted CD8 T cells. On the other hand, GLUT2, which has been reported to have low affinity to glucose, can be induced during T cell activation, peaking earlier than GLUT1, and is crucial for maintaining nutrient uptake and metabolism in activated CD8 T cells rather than naïve T cells or activated CD4 T cells [46]. Therefore, the functions of GLUT family members on CD8 cells may deserve more experimental digging to sort out their potential roles.

Increased glucose uptake and aerobic glycolysis are primary features of cancer cells as well [47]. Aerobic glycolysis is a process in which glucose is converted into pyruvate, which eventually becomes lactate, with a small amount of ATP produced [48]. Excessive division, invasion and metastases of malignant tumor cells alter their energy demands from aerobic respiration to glycolysis, resulting in enhanced production of glycolytic enzymes and glucose transporter proteins [49]. Oncogenic transformation in cultured tumor cells causes strong glucose transport and increased expression of GLUT1 and GLUT3 [50]. Positive correlations between glucose uptake and levels of GLUT1, GLUT3 or GLUT12 were observed in many different types of cancers that are associated with poor survival rates [51]. These findings indicate that any anti-tumor therapy targeting glycolysis in the tumor microenvironment is a double-edged sword requiring enhancement of the glycolytic pathway to favor effector CD8 T cells while excluding side effects that promote the aggressive progression of tumor cells. BCL6 directly represses genes involved in the glycolysis pathway, including *Slc2a1* and *Slc2a3* [52]. Although, BCL6 associates with the promoters of many genes of this family [53], there is little functional information available about other members. Increasing studies suggest that BCL6 works as an oncogene in human cancer [54]. BCL6 is a powerful transcriptional repressor that silences a variety of genes involved in the DNA damage response, cell cycle and death, such as TP53, CDKN1A/B, ATR, CHEK1, CDKN2A/B, PTEN, etc. [55]. In the present study, the antitumor efficacy of Fx1 relied on immune cells, suggesting BCL6 expression in MC38 or B16F10 tumor cells likely did not reflect a direct anti-tumor effect, though inhibiting BCL6 in tumors may benefit tumor growth resistance as well. Ultimately, our data introduce an effective therapeutic strategy by targeting effector CD8 T cells to improve their glycolysis, thereby prolonging their antitumor cytotoxic functions.

## Acknowledgments

This study was supported by National Institute of Health (R01AI137252 to C.X. and D.N.). We thank members of Xiao and Nemazee laboratories for their discussion.

## Author Contributions

F.L. and Y.L. designed and performed the experiments, analyzed data, and contributed to the writing of the manuscript; N.J. performed the cell mitochondria stress tests. J.T.T, T.R.B. and R.B. provided mice for experiments. Y.L., Z.H., C.X., and D.N. conceived and designed this study. The initial discovery was made in the Xiao lab, and the study was continued in the Nemazee lab due to the departure of C.X. from The Scripps Research Institute in 2021. Y.L. and D.N. wrote the manuscript with contributions from other authors.

---

**Figure S1.**
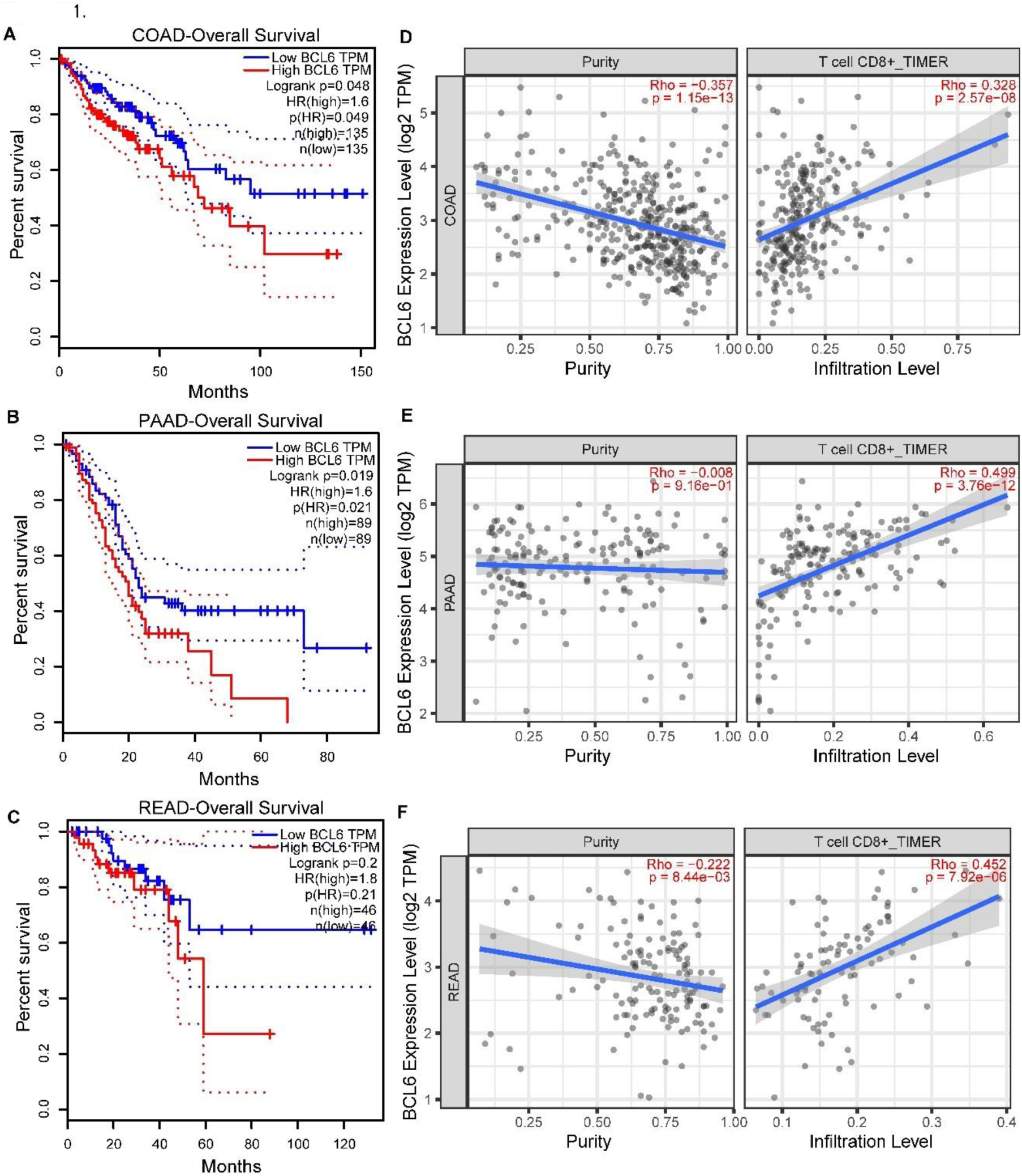
Relationship between BCL6 expression level, overall survival rate and CD8+ T cell infiltration in tumors from TIMER2.0 database. **(A-F)** In COAD (A, D), PAAD (B, E) and READ (C, F) patients, BCL6 expression and overall survival exhibited a negative correlation but a positive correlation was found between BCL6 levels and CD8 T cell infiltration.

**Figure S2.**
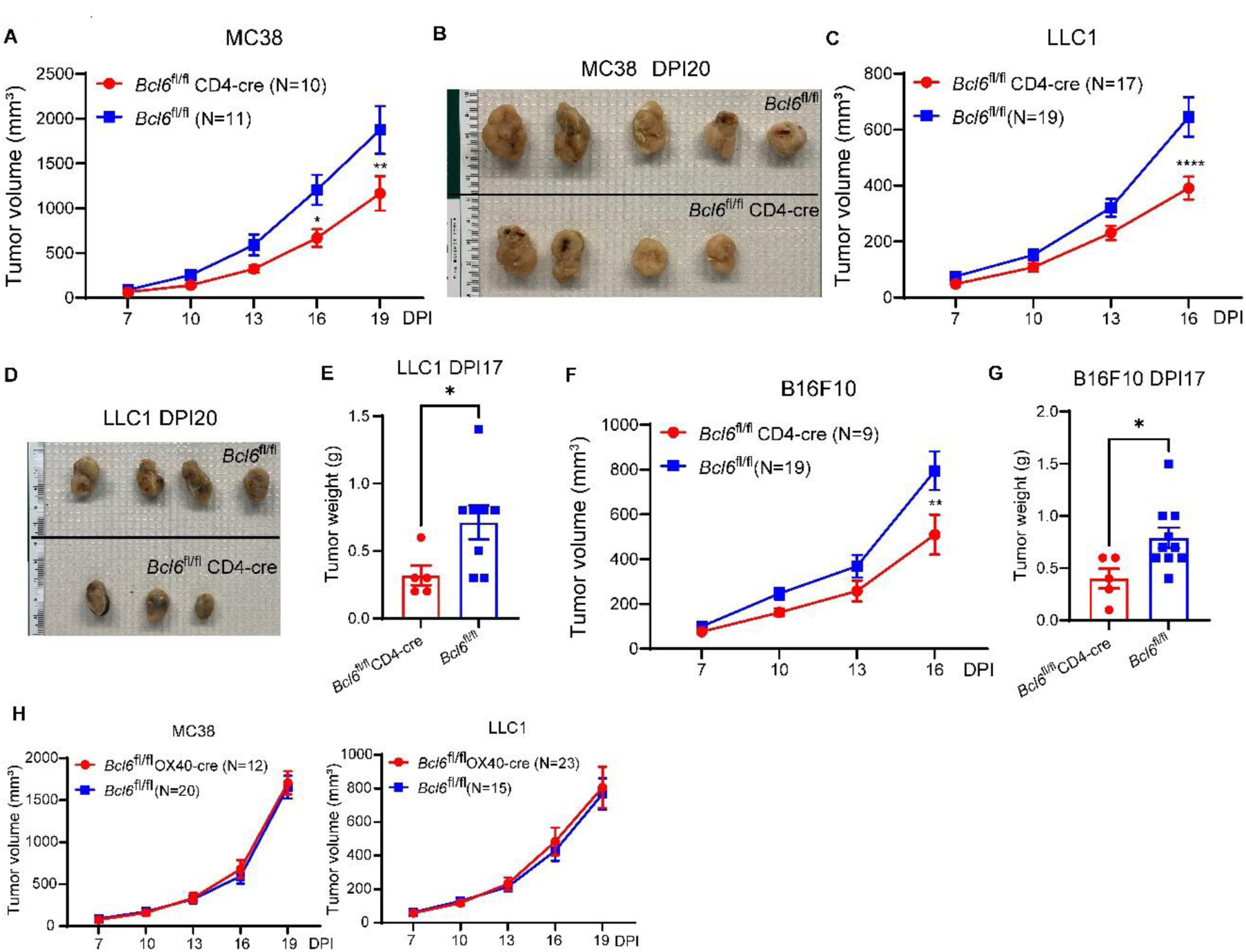
Analysis of tumor growth in *Bcl6*^fl/fl^ CD4-cre and *Bcl6*^fl/fl^ OX40-cre mice compared to control *Bcl6*^fl/fl^ mice. **(A-G)** Compared to *Bcl6*^fl/fl^ control mice, growth of MC38 (A and B), LLC1 (C-E), B16F10 (F-G) were significantly repressed in *Bcl6*^fl/fl^ CD4-cre mice. Tumor growth curves shown in A, C and F; photos in B and D were taken on DPI20; and tumor weights in E and G were assessed on DPI 17. Tumor inoculation data were from 2-3 independent experiments. (H) No difference in progression of MC38 (left) or LLC1 (right) was found comparing *Bcl6*^fl/fl^ OX40-cre and *Bcl6*^fl/fl^ control mice. Data were pooled from 3-4 independent experiments. Each dot in column graphs (E-G) represents an individual mouse. Bars show mean ± SEM. P values were calculated with the two-tailed unpaired t-test (E-G), Two-way ANOVA (A, C-F). *, **, **** denote P < 0.05, 0.01, 0.0001, respectively.

**Legend of Supplementary Dataset 1 (Excel file).** Significantly differentially expressed genes between BCL6-knockout (BCL6-KO) and BCL6-WT activated CD8 T cell samples n=4 per genotype group, padj < .05.

